# Disruption of *fos* causes craniofacial anomalies in developing zebrafish

**DOI:** 10.1101/2022.12.06.519379

**Authors:** Lorena Maili, Bhavna Tandon, Qiuping Yuan, Simone Menezes, S. Shahrukh Hashmi, Ariadne Letra, George T. Eisenhoffer, Jacqueline T. Hecht

## Abstract

Craniofacial development is a complex and tightly regulated process and disruptions can lead to structural birth defects, the most common being nonsyndromic cleft lip and palate (NSCLP). Previously, we identified *FOS* as a candidate regulator of NSCLP through family-based association studies, yet its specific contributions to oral and palatal formation are poorly understood. This study investigated the role of *fos* during zebrafish craniofacial development through genetic disruption and knockdown approaches. *Fos* was expressed in the periderm, olfactory epithelium and other cell populations in the head. Genetic perturbation of *fos* produced an abnormal craniofacial phenotype with a hypoplastic oral cavity that showed significant changes in midface dimensions by quantitative facial morphometric analysis. Loss and knockdown of *fos* caused increased cell apoptosis in the head, followed by a significant reduction in cranial neural crest cells (CNCCs) populating the upper and lower jaws. These changes resulted in abnormalities of cartilage, bone and pharyngeal teeth formation. Periderm cells surrounding the oral cavity showed altered morphology and a subset of cells in the upper and lower lip showed disrupted Wnt/β-catenin activation, consistent with modified inductive interactions between mesenchymal and epithelial cells. Taken together, these findings demonstrate that perturbation of *fos* has detrimental effects on oral epithelial and CNCC-derived tissues suggesting that it plays a critical role in zebrafish craniofacial development and a potential role in NSCLP.

**Summary statement:** Perturbation of *fos*, a candidate gene associated with nonsyndromic cleft lip and palate in humans, causes a distinctive orofacial phenotype in zebrafish as a result of abnormal development of craniofacial tissues. Disruption of *fos* in the oral epithelial, cranial neural crest and Wnt responsive cell populations around the oral cavity causes anomalies that suggest a potential role in the etiology of NSCLP.

## INTRODUCTION

Vertebrate craniofacial development relies on a complex and tightly regulated series of tissue growth and fusion, and even minor disruptions to this intricate process can lead to birth defects, the most common of which is nonsyndromic cleft lip and palate (NSCLP) (Cordero et al., 2011; Gorlin, 2011). The cranial neural crest cells (CNCCs) are a multipotent population of cells that give rise to the various tissues comprising the craniofacial structures, including cartilage and bone, muscles, sensory neurons, and pigment cells (Cordero et al., 2011; Dash & Trainor, 2020; Mork & Crump, 2015). In human embryonic development, CNCCs give rise to the facial prominences during the fourth week of gestation, which over the subsequent 6-8 weeks, grow, make contact and fuse to form complete facial structures (Marazita & Mooney, 2004). Additionally, epithelial cells in the facial prominences provide important signals to CNCCs, directing their proliferation and patterning to further regulate craniofacial growth and morphogenesis (Chai & Maxson, 2006; Jiang, Bush, & Lidral, 2006). These interactions involve multiple molecular signaling pathways, including, but not limited to, wingless-type MMTV integration site family (WNT), fibroblast growth factor (FGF), bone morphogenetic protein (BMP) and ectodysplasin A (EDA) (Chai & Maxson, 2006; Reynolds et al., 2019).

Deficiencies in cell migration, changes in mitotic activity or apoptosis, and interruptions to epithelial-mesenchymal interactions can affect the shape and size of the facial prominences and increase the likelihood of an orofacial cleft (Dixon, Marazita, Beaty, & Murray, 2011; Ji et al., 2020; Jiang et al., 2006). Quantitative studies of facial form using geometric morphometrics (GMM) have provided important information for understanding processes disrupted during craniofacial development by identifying phenotypic changes in facial prominences that contribute to orofacial clefts (Young, Wat, Diewert, Browder, & Hallgrimsson, 2007).

In previous studies, we reported an association of *CRISPLD2* with NSCLP that was independently replicated in different populations (Chiquet et al., 2007; Ge et al., 2018; Letra et al., 2010; Mijiti et al., 2015; Shen et al., 2011). Further, we showed that knockdown and loss of *crispld2* in zebrafish caused severe craniofacial defects resulting from altered migration of CNCCs (Chiquet et al., 2018; Swindell et al., 2015; Yuan et al., 2012). RNA-seq analysis of *crispld2* wild type (WT) and morphant embryos revealed that *fosab* (zebrafish homologue of *FOS*) was differentially expressed. This finding, and a positive association between *FOS*/rs1046117 and NSCLP, identified *FOS* as a novel candidate NSCLP gene (Chiquet et al., 2018).

*Fos* is a transcription factor as part of the activating protein-1 (AP-1) transcription factor complex (Durchdewald, Angel, & Hess, 2009). *FOS* has roles in oncogenic processes such as tumor growth and progression, as well as biological processes like proliferation, differentiation, epithelial-to-mesenchymal transition and apoptosis (Chen et al., 2017; Durchdewald et al., 2009; Milde-Langosch, 2005; Rodriguez-Berdini et al., 2020; Velazquez et al., 2015; Wagner, 2002). Fos is also an activator in the lipid synthesis pathway in the endoplasmic reticulum (Durchdewald et al., 2009; Rodriguez-Berdini et al., 2020; Wagner, 2002). Mice lacking *Fos* display severe osteopetrosis, impaired gametogenesis, abnormal hematopoiesis and behavioral changes (Grigoriadis, Wang, & Wagner, 1995). Craniofacial abnormalities such as a domed skull with a shorter snout, absence of tooth eruption, and a reduced neocortex are also common features in *Fos* null mice (Alfaqeeh et al., 2015; Johnson, Spiegelman, & Papaioannou, 1992; Velazquez et al., 2015; Wagner, 2002). Importantly, reporter expression is found in orofacial tissues, including the medial edge epithelium (MEE) of the palate, the dental papilla mesenchyme, the periderm, and cells at the midline of the nasal septum of *fos-lacZ* mice (Smeyne, Schilling, et al., 1993; Smeyne, Vendrell, et al., 1993). Fos protein is also detected in the rat MEE just before the elevation of the palatal shelves, the facial epidermis, Meckel’s cartilage and the mesenchymal condensations that precede bone and muscle formation (Yano, Ohtsuru, Ito, Fujii, & Yamashita, 1996). These studies provide strong evidence that *FOS* plays a role during craniofacial development. However, its contributions to mouth and palate formation remain to be elucidated. This study investigated the role of *fos* during craniofacial development in zebrafish using geometric morphometrics combined with tissue and cell analysis to determine its potential involvement in craniofacial morphogenesis and potential contributions to NSCLP.

## RESULTS

### *Fos* is expressed in midfacial and perioral regions during zebrafish development

Fluorescent *in situ* hybridization via hybridization chain reaction (HCR)(Choi et al., 2018) was used to evaluate *fosab* (referred to as *fos*) mRNA expression during 1 to 5 days post fertilization (dpf) of zebrafish development. Overall, low levels of *fos* mRNA were observed in the embryonic head at 1 dpf, however, expression was detected in a subset of tissues around the eyes (**Fig.1 A, A’, B, B’; Supplemental Figure 1A**). Increased *fos* expression was observed at 2 dpf as well as around the nares/olfactory placodes, midfacial region, eyes, oral cavity and brain (**Fig. 1B, 1B’ 1C, 1C’**). By 5dpf, strong expression was observed in the areas surrounding the nares/olfactory pits, perioral tissues (upper and lower lips), and more diffusely in the lower jaw (**Fig 1 D, 1D’, 1E, 1E’**). Expression was also observed in the otic vesicle and lens (**Fig. 1D, 1D’**). Examination of individual z-slices at 5 dpf showed fluorescent signal in the brain and olfactory epithelium, and the outer epithelial layer (**Supplemental Fig. 2**). Quantitative real-time PCR of *fos* transcripts at these developmental time points supported the *in situ* results, with lower *fos* expression observed at 1 dpf and highest at 5 dpf (**Fig. 1H**). These findings support the expression of *fos* in craniofacial tissues throughout early development stages in zebrafish.

**Figure 1.**
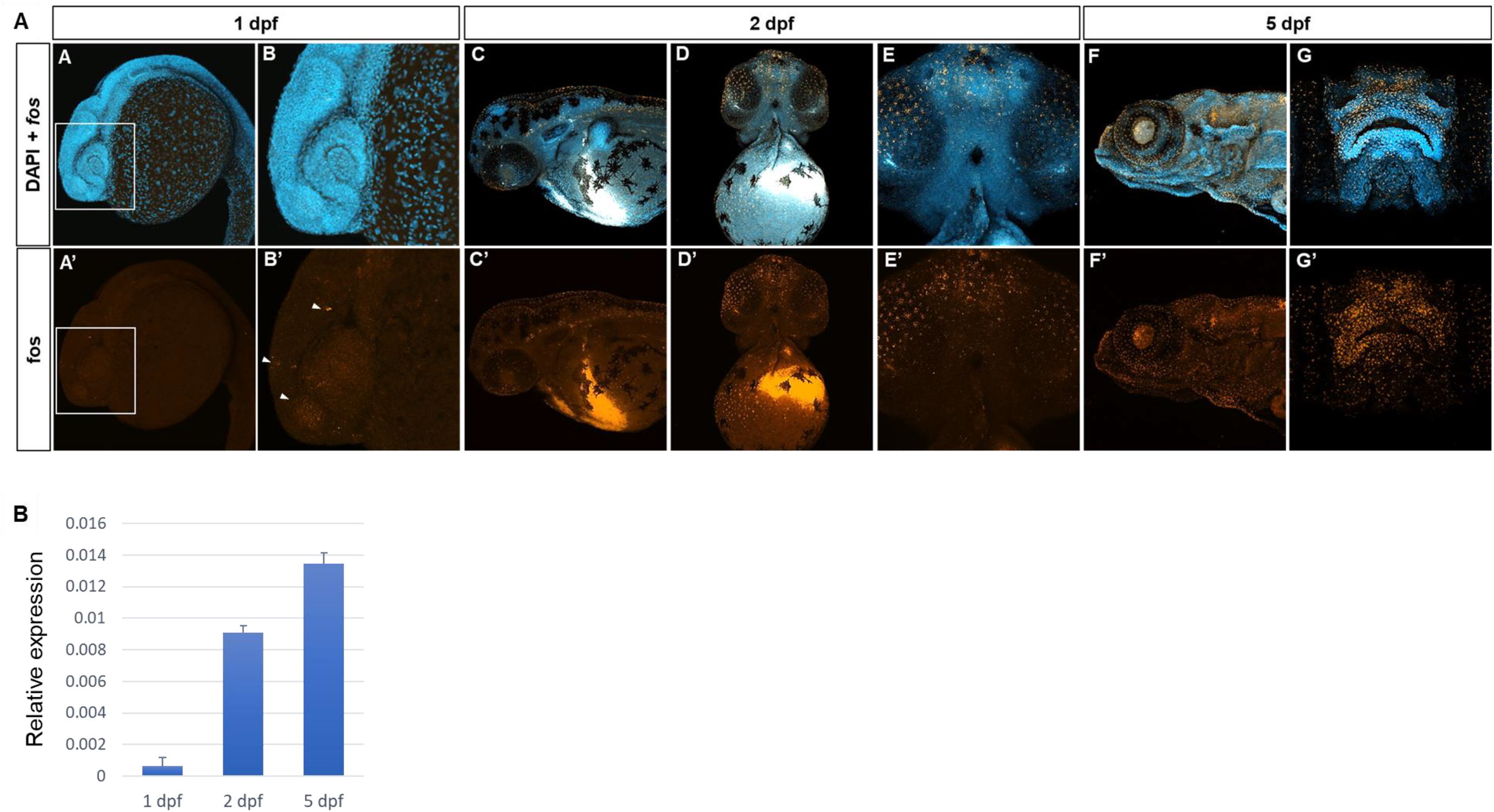
*Fos* is expressed in craniofacial tissues during zebrafish development. **Panel A)** In situ hybridization with zebrafish *fosab* RNA probe (HCR assay) showed localized expression around the developing brain and olfactory placodes at 1 dpf, shown in lateral view and denoted by arrowheads (**A, A’, B, B’),** increased expression in the midface and oral cavity at 2 dpf shown in lateral and ventral views (**C, C’, D, D’, E, E’)** and 5 dpf show in lateral and rostral views (F, F’, G, G’). **Panel B)** Quantitative RT-PCR showing relatively lower mRNA expression at 1 dpf and higher expression at 2 dpf and 5 dpf.

### *Fos* perturbation causes an abnormal phenotype in zebrafish

To define the role of *fos* in craniofacial development, zebrafish F0 mutants (referred to as crispants (Burger et al., 2016)) were generated using CRISPR/Cas9 with two single guide RNAs (sgRNAs) targeting exons 1 and 4 of the *fos* gene. Genotyping revealed efficient mutagenesis with 89% of embryos showing a deletion of approximately 1500 base pairs (bp) of the *fos* coding region (**Fig. 2A**). F0 embryos (84%) displayed an abnormal phenotype at 5 dpf, with smaller head and eyes, cardiac edema, abnormal yolk extension, missing swim bladder and curved body axis compared to uninjected control embryos (UIC) and tyrosinase sgRNA-injected F0 embryos (0%) (**Fig. 2B, C. Supplemental Fig. 4**). Abnormal mouth shape, displaced neuromasts, and anomalies in lower jaw tissues were observed (**Fig 2D, E**). *In situ* hybridization showed reduced *fos* mRNA expression in the head at all developmental time points examined (**Supplemental Fig. 1**). A small number of embryos (16%) appeared normal, likely reflecting mosaicism resulting from F0 CRISPR mutagenesis. Stable *fos* mutants (F2 generation) with the same deletion as F0 crispants had significantly reduced *fos* mRNA levels and a normal phenotype. Genetic compensation studies of closely related genes showed that *fosb* was significantly upregulated in the mutants at both 2 and 3 dpf (p = 0.03 and 0.006 respectively) (**Supplemental Fig. 3**). Moreover, injection of the *fos* guide sgRNAs into stable F2 mutants did not cause an abnormal craniofacial phenotype (**Supplemental Fig. 4A**).

**Figure 2.**
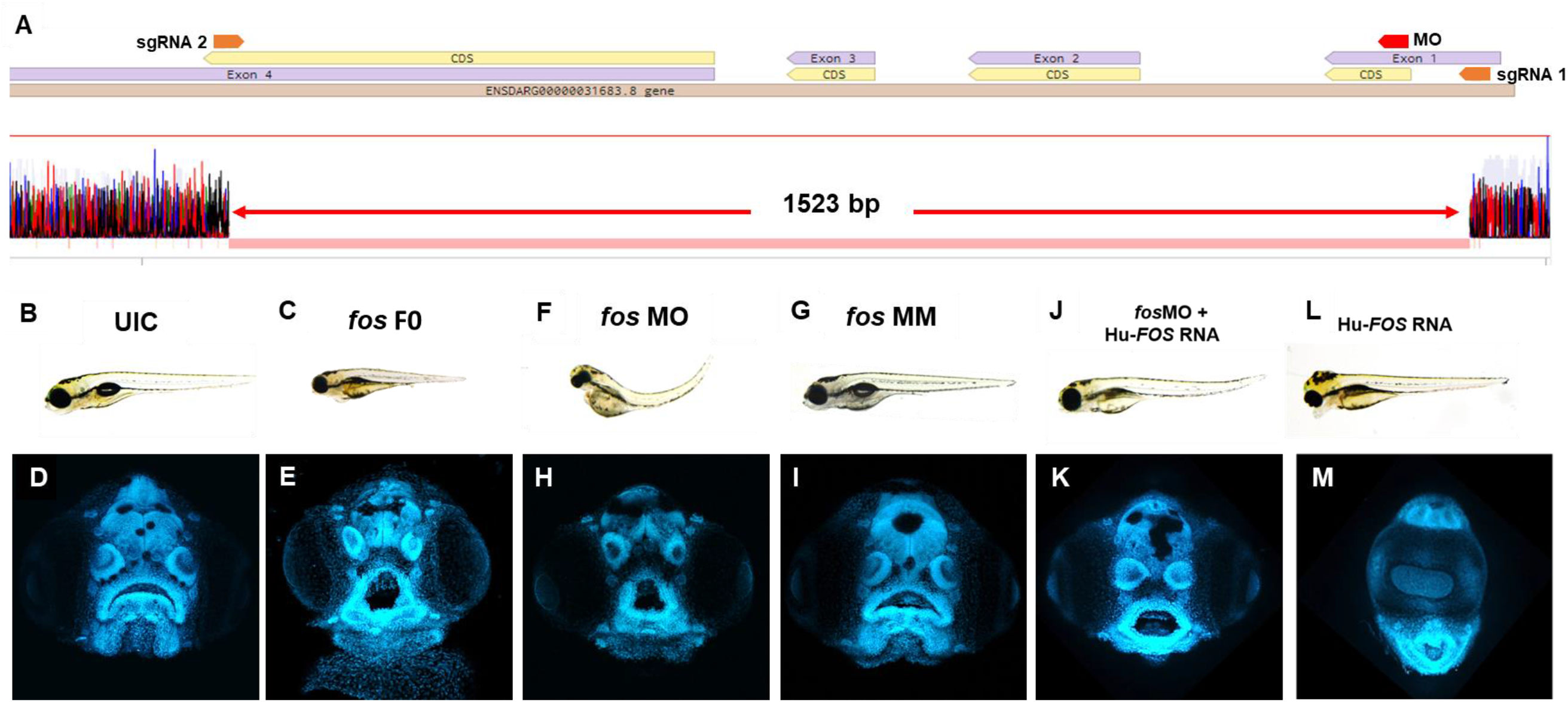
*Fos* mutants and morphants have distinctive keyhole-shaped mouth and abnormal craniofacies. **A)** Schematic of CRISPR guides targeting exons 1 and 4 of the *fosab* gene and resulting in an almost complete deletion of the coding region with a 1.523 kb deletion confirmed by DNA sequencing. **B-L)** Representative brightfield images of 5dpf zebrafish showing normal phenotype (B) and abnormalities in *fos* F0 mutants and morphants (C, F) compared to UIC and MM controls (B, G). **D-M)** Confocal images of rostrally mounted DAPI-stained samples showed a distinctively abnormal facial and mouth shape in both *fos* mutant (H) and morphant (I) embryos compared to UIC (D) and MM controls (I). Co-injection of human *FOS* mRNA rescued the morphant phenotype (J, K) while overexpression alone caused other abnormalities including cyclopia and an elongated lower jaw (L, M). UIC, uninjected control; MO, morpholino; MM, mismatch; Hu-FOS, human *FOS* mRNA.

To further evaluate the consequences of reduced *fos* on craniofacial development, a translation-blocking MO against *fos* was used. *Fos* morphant embryos displayed similar abnormalities to crispants at 5 dpf (**Fig. 2F, H**), while mismatch (MM) control injected embryos were similar to the UICs (**Fig. 2G, I**). Co-injection with a morpholino directed to p53 did not improve the phenotype, supporting that the abnormal morphant phenotype was independent of non-specific cell death (ref) (**Supplemental Fig. 4B**). To test the specificity of *fos* knockdown, human *FOS* (Hu-FOS) mRNA not targeted by the *fos* MO was co-injected and rescued the morphant abnormalities including the size of the head and eyes, as well as oral cavity shape (**Fig. 2J, K**). While 82% of *fos* morphants had an abnormal mouth and face, only 45% had an abnormal phenotype after human FOS mRNA co-injection (**Supplemental Fig. 5A**). Interestingly, overexpression of human-*FOS* RNA in zebrafish embryos caused severe facial abnormalities, including an extended lower jaw and cyclopia (**Fig. 2L, M**). Pharmacological perturbation of AP-1 transcription factor complex activity mediated by *fos* using SR 11302 resulted in craniofacial abnormalities similar to those observed in *fos* crispant and morphant embryos, including narrow face, smaller mouth, and altered olfactory pits (**Supplemental Fig. 6**). This resulting phenotype was more severe, as expected because of the wider range of disruption by the AP-1 inhibitor, which targets the AP-1 dimer complex composed of different members of the fos and jun protein families (*fos*, *fosb, fra-1, fra-2, jun, junB, junD*) (Wagner, 2002). Taken together, these data suggest a key role for *fos* in regulating craniofacial development in zebrafish embryos.

### Geometric morphometric analysis identifies orofacial features altered by *fos*

To identify the orofacial features altered by genetic perturbation of *fos* at 5 dpf, we used zFACE, a geometric morphometric tool for quantitative deep phenotyping of the craniofacial region in zebrafish embryos (Maili et al., 2022). This tool uses 26 anatomical landmarks, comparable to those used in human geometric morphometric (GMM) studies (Weinberg et al., 2008), to calculate 39 facial measurements, including linear distances, areas and angles (Maili et al., 2022). As shown in **Table 1**, significant alterations were observed for 16 facial features in both *fos* crispants and morphants compared to UIC and MM controls (ANOVA with post-hoc Tukey’s test, p < 0.0001). These included differences in oral cavity dimensions and angles, olfactory placode positioning, as well as area measurements in the lower half of the face. Similar phenotypic results were observed at 6 dpf, indicating that the facial anomalies did not improve with time and were not the result of a delay in development (**Supplemental Fig. 7, Supplemental Table 1**). Nine of the 16 altered dimensions (2 olfactory to mouth angles, mouth height, and 6 oral cavity angles) were fully rescued by Hu-FOS mRNA injection (**Table 1, Figure 3A**). Olfactory distance, neuromast width and mouth perimeter were partially rescued; they were different compared to the controls and to the morphants, representing an intermediate phenotype (**Table 1, Fig. 3B**). Four facial features, chin width, a neuromast angle, area bottom and area combined, were not rescued (**Table 1, Figure 3C**). Morphometric analysis could not be performed in the overexpression group due to the severity of the phenotype and many anatomical landmarks missing.

**Figure 3.**
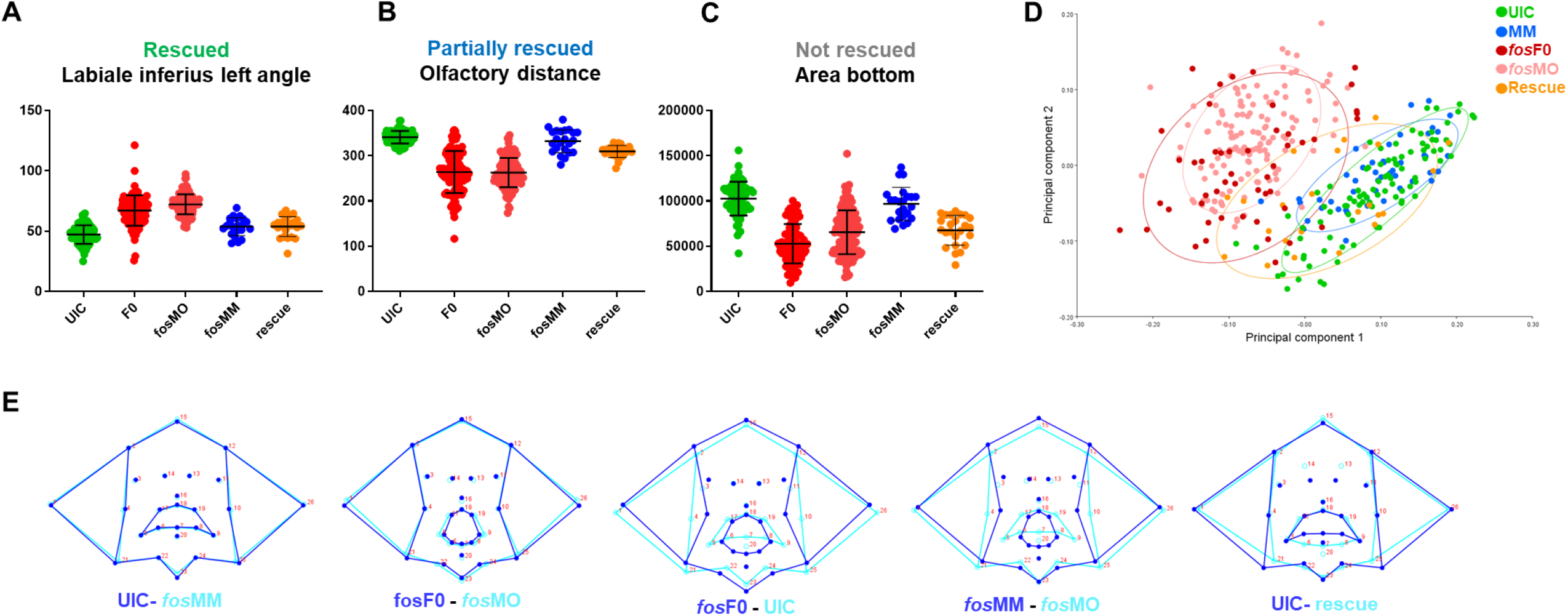
Quantitative analysis confirms facial abnormalities in *fos* crispants and morphants. **A)** ZFACE morphometrics identified significant measurement differences in 16 individual craniofacial parameters for *fos* mutants and morphants compared to MM/UIC controls. Nine of these parameters were completely rescued (only the labiale superius left, an oral cavity angle, is shown as an example). **B)** Three parameters were partially rescued (olfactory distance shown as an example) while **C)** 4 were not rescued (area of the bottom half of the face shown). **D)** Principal component analysis of Procrustes transformed landmarks showed an overlap of *fos* crispant and morphant groups, as well as separation from the UIC and MM groups, while the rescue group plotted closer to the controls. **E**) Summary of overall shape differences between control and experimental groups, highlighting the similarities between UIC and MM controls, crispants and morphants, and the differences between UIC/MM controls and crispants/morphants. Additionally, rescue with *fos* mRNA showed a more normal average mouth and face shape.

**Table 1.** Summary of morphometric analysis results.

| zFACE feature | ANOVA | fos F0 mutants |  | fos morphants |  |  | rescue |  | Summary |
| --- | --- | --- | --- | --- | --- | --- | --- | --- | --- |
|  |  | UIC vs. F0 | UIC vs. fosMO | UIC vs. fosMM | fosMM vs. fosMO | fosMO vs. F0 | UIC vs. rescue | fosMO vs. rescue |  |
| Olfactory to Mouth 2 | <0.0001 | <0.0001 | <0.0001 | n.s. | 0.0003 | n.s. | n.s. | <0.0001 | rescued |
| Olfactory to Mouth 3 | <0.0001 | <0.0001 | <0.0001 | n.s. | 0.0002 | n.s. | n.s. | <0.0001 | rescued |
| Mouth Height | <0.0001 | <0.0001 | <0.0001 | n.s. | <0.0001 | n.s. | n.s. | <0.0001 | rescued |
| Labiale Inferius Angle | <0.0001 | <0.0001 | <0.0001 | n.s. | <0.0001 | 0.008 | n.s. | <0.0001 | rescued |
| Crista Philtri Left Angle | <0.0001 | <0.0001 | <0.0001 | n.s. | <0.0001 | 0.005 | 0.002 | <0.0001 | rescued |
| Crista Philtri Right Angle | <0.0001 | <0.0001 | <0.0001 | n.s. | <0.0001 | n.s. | 0.005 | <0.0001 | rescued |
| Labiale Superius Mid Angle | <0.0001 | <0.0001 | <0.0001 | n.s. | <0.0001 | 0.003 | n.s. | <0.0001 | rescued |
| Labiale Inferius Left Angle | <0.0001 | <0.0001 | <0.0001 | n.s. | <0.0001 | n.s. | n.s. | <0.0001 | rescued |
| Labiale Inferius Right Angle | <0.0001 | <0.0001 | <0.0001 | n.s. | <0.0001 | n.s. | n.s. | <0.0001 | rescued |
| Olfactory Distance | <0.0001 | <0.0001 | <0.0001 | n.s. | <0.0001 | n.s. | <0.0001 | <0.0001 | partial rescue |
| Neuromast Width | <0.0001 | <0.0001 | <0.0001 | n.s. | <0.0001 | 0.002 | <0.0001 | 0.002 | partial rescue |
| Mouth Perimeter | <0.0001 | <0.0001 | <0.0001 | n.s. | <0.0001 | n.s. | <0.0001 | <0.0001 | partial rescue |
| Chin Width | <0.0001 | <0.0001 | <0.0001 | n.s. | <0.0001 | n.s. | 0.003 | n.s. | not rescued |
| Neuromast Angle 2 | <0.0001 | <0.0001 | <0.0001 | n.s. | <0.0001 | n.s. | 0.002 | n.s. | not rescued |
| Area Bottom | <0.0001 | <0.0001 | <0.0001 | n.s. | <0.0001 | n.s. | <0.0001 | n.s. | not rescued |
| Area Combined | <0.0001 | <0.0001 | <0.0001 | n.s. | <0.0001 | 0.005 | 0.0002 | n.s. | not rescued |

Unbiased multivariate analysis using principal component analysis (PCA) was next performed after Procrustes transformation to determine overall shape changes after *fos* disruption. The first two components (PC1, PC2) were responsible for 63% of the variance in the dataset. Landmarks in the lower part of the oral cavity and midface had the largest vectors of change across PC1, while landmarks outlining the lower jaw, location of the pupils and upper lip had the largest vectors of change across PC2 (**Supplemental Fig. 8**). A component plot for PC1 and PC2 showed overlap of *fos* crispant and MO groups, as well as separation of the crispant and MO groups from the UIC and MM controls (**Fig. 3D**). A small subset (n = 5, 10%) of crispants clustered close to the UIC and MM control groups, likely due to mosaicism. The rescue group mapped closer to the control groups, with some individuals in the intermediate area of the plot, likely reflecting the partial rescue observed in the previous analysis (**Fig. 3D**).

Shape analysis was also performed using discriminant function analysis (DFA) to determine face shape differences between experimental and control groups (Klingenberg, 2011). As expected, face shape in UIC and MM controls was similar to each other (Procrustes distance = 0.03, p = 0.06). Although a significant difference was found between *fos* crispants and morphants, comparison of wireframe representations revealed only minor alterations in face shape (Procrustes distance = 0.06, p < 0.001) (**Fig. 3E**). Comparisons between both crispants to UICs (Procrustes distance = 0.18, p < 0.001) and morphants to MM controls (Procrustes distance = 0.17, p < 0.001) showed significant shape differences involving the oral cavity, midface and lower jaw in both comparisons (**Fig. 3E**). DFA also highlighted the improvement in the rescue group when compared to UICs, specifically in the upper lip and upper face regions, however the lower lip and jaw landmarks remained abnormal (**Fig. 3E**). Together, these quantitative analyses demonstrated that perturbation of *fos* affected oral cavity and midface dimensions and overall facial shape.

### *Fos* perturbation alters craniofacial structures

Given that the shape of the underlying skeletal tissues is a major contributing factor to facial shape, the bone and cartilage based structures that comprise craniofacial tissues were examined (Murillo-Rincon & Kaucka, 2020). Skeletal staining showed reduced and abnormally shaped jaw cartilages, including smaller ethmoid plate, trabeculae and parachordal cartilages, reduced Meckel’s and palatoquadrate cartilages and missing basibranchial cartilages in both *fos* crispants and morphants (**Fig. 4 B, B’, C, C’**) compared to UIC and *fos* MM controls (**Fig. 4 A, A’, D, D’**). The parasphenoid bone, branchiostegal rays and fifth ceratobranchial arches (CB5) showed impaired ossification and smaller or missing pharyngeal teeth (**Fig.4 B-C**, asterisks). Fused occipital bones and asymmetric neurocranial structures were also observed (**Fig 4C’**). These abnormalities were dose-dependent in the knockdown experiments, ranging from mild (0.5ng/uL MO injection) to very severe (2ng/uL MO injection) (**Supplemental Fig. 9**).

**Figure 4.**
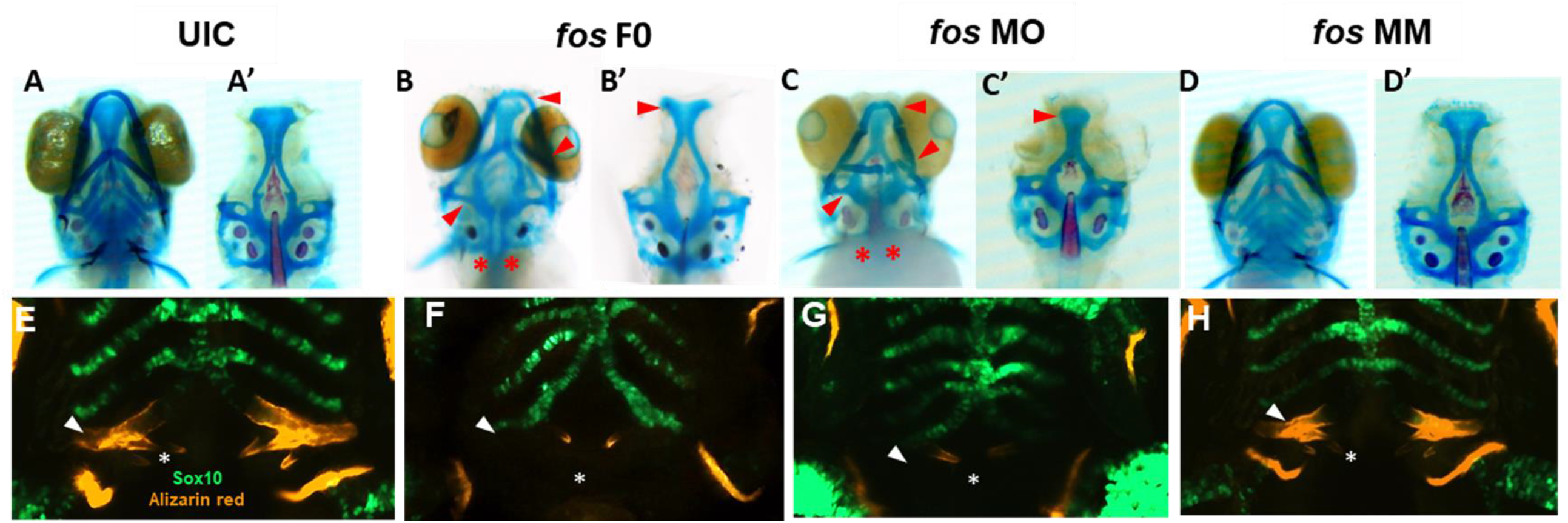
Perturbation of *fos* causes bone, cartilage, and tooth abnormalities. **A-D)** *Fos* crispants and morphants show abnormally shaped jaw cartilages, including a smaller ethmoid plate, Meckel’s and palatoquadrate cartilages, missing basibranchial cartilages (B – C’ red arrows) and missing pharyngeal teeth (red asterisks). **E-H)** Only 24% of *fos* crispant and morphant zebrafish showed mineralization at the fifth ceratobranchial arch (F, G, white arrows) compared to 100% of the UIC/MM (E, H). Tooth number was reduced and tooth shape/size was altered in *fos* crispants and morphants compared to UIC/MM (**E-H**, white asterisks).

The multiple bone and cartilage anomalies suggested abnormal pharyngeal/branchial arch formation. *Fos* knockdown in the *Tg(−4.9sox10:GFP)* zebrafish line was used to visualize the developing pharyngeal arches (PAs) (Carney et al., 2006). At 5dpf, PAs 1 and 2 were abnormal, with crispants/morphants showing cellular disorganization, vertical and horizontal constriction and arch asymmetry (**Supplemental Fig. 10**). Approximately 20% of crispant, and 37% of morphant larvae showed merged PA1 and PA2; while all crispants/morphants had abnormally-shaped but intact PAs 3-7 (**Fig. 4 G-H** and **Supplemental Fig. 10**). Additionally, mineralization of the fifth ceratobranchial (CB5/PA 7), visualized by alizarin red staining, was observed in 100% of the UIC and MM embryos, compared to only 19% of *fos* crispants and 24% of morphants (**Fig. 4E-H** arrow, **Supplemental Fig. 11, Supplemental Table 2**). Decreased mineralization persisted during subsequent developmental stages, with only 68% of morphants showing evidence of mineralization by 8dpf (**Supplemental Fig. 11D**). Smaller cleithrums and opercles were also observed, indicating abnormal ossification in craniofacial dermal bones (**Fig. 4 E-H**).

Development of zebrafish pharyngeal dentition was also examined in *fos* crispants and morphants. At 5dpf, 3 bilateral rows of pharyngeal teeth (4V1, 3V1 and 5V1), were observed in both UIC and MM controls, with 4V1 ankylosed/attached to the perichondral bone of CB5, as expected (**Fig. 4, E, H; Supplemental Fig. 11C**). In comparison, crispant and morphant larvae had only 1 tooth (4V1) on each side of the arch, which was smaller and of abnormal morphology compared to the controls. In mild phenotypic crispants/morphants, 4V1 had attached/deposited bone around the non-mineralized CB5 cartilage, while in crispants/morphants showing severe phenotypes only 4V1 buds were visible (**Fig. 4F, G; Supplemental Fig. 11C**). Defects in tooth attachment, morphogenesis, and in CB5 mineralization persisted to 8dpf (**Supplemental Fig. 11D, Supplemental Table 2**). These abnormalities in bone, cartilage, pharyngeal arch and tooth formation in both *fos* crispants and morphants are consistent with perturbation of a common developmental program.

### Epithelial and neural crest cell alterations result from *fos* disruption

Cell populations important for craniofacial development in *fos* crispants and morphants were next examined to understand the abnormal phenotype resulting from *fos* perturbation. Since expression of *fos* mRNA was detected in the outer epithelial layer, epithelial cell number and morphology were analyzed using a keratin-4 transgenic reporter line *Tg(krt4:gfp)* (Gong et al., 2002). *Fos* crispants and morphants showed abnormal cellular arrangement, particularly around the oral cavity and nares/olfactory pits compared to UIC and MM controls at 3 and 5 dpf (**Fig. 5 A-D**). Visualization of the borders and cell-cell junctions of epithelial cells also further showed the smaller oral cavities and olfactory pits in crispants/morphants (**Fig. 5 E-H**). In addition to altered arrangement of epithelial cells around the oral cavity and lower jaw, cells in the perioral region of both crispants and morphants had a smaller area compared to controls at both 3 dpf (p < 0.0001, Student’s t-test) and 5 dpf (p < 0.0001) (**Fig. 5 I, J**; see **Supplemental Fig. 12** for a description of facial region analyses). These results suggested a role for *fos* in regulating epithelial cell size and arrangement around the oral cavity.

**Figure 5.**
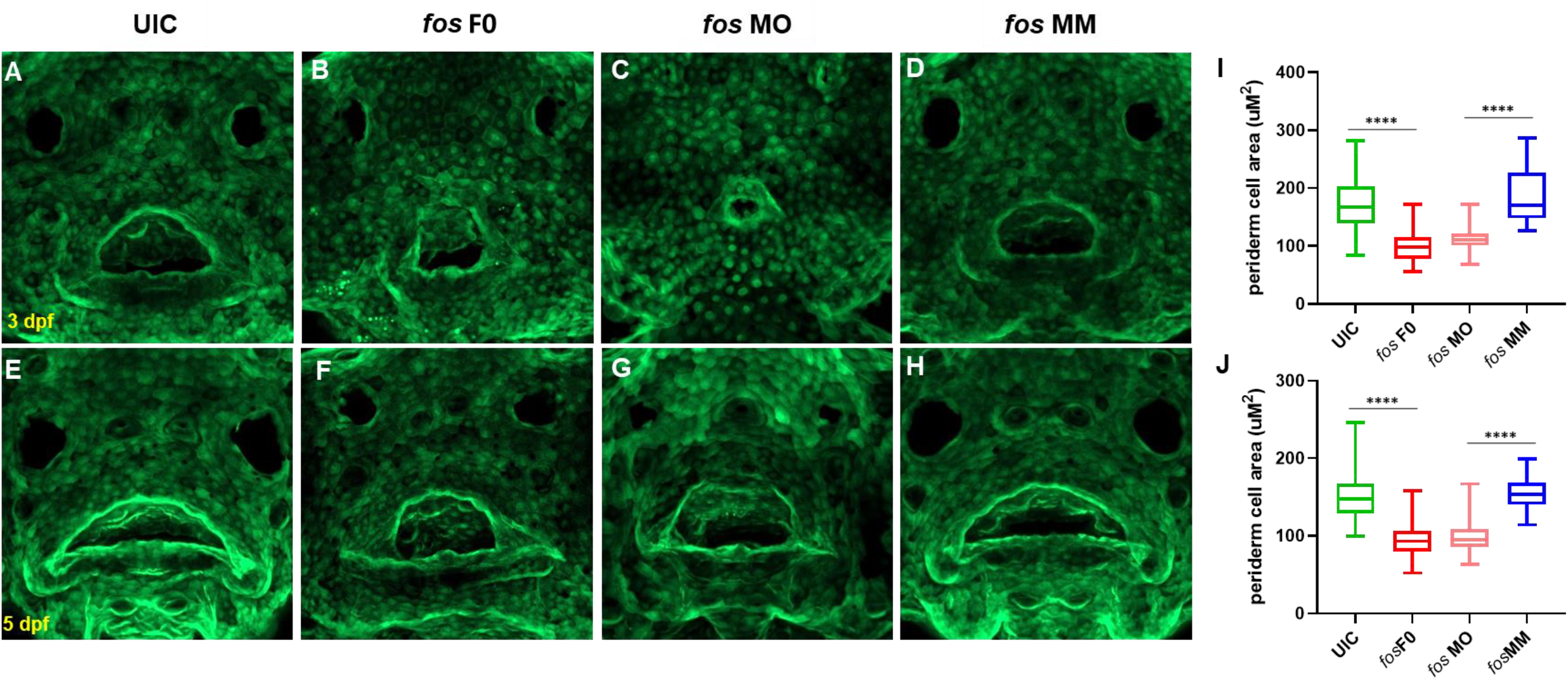
Perturbation of *fos* causes epithelial abnormalities. **A-H**) Examination of Keratin-4 expressing cells showed abnormal arrangements of peridermal epithelial cells around the oral cavity in both crispants (**B, F**) and morphants (**C, G**) compared to controls at both 3dpf (**A, D**) and 5dpf (**E, H**). Periderm cell size was quantified and both crispants and morphants (minimum of n = 60 cells per condition) showed reduced perioral cell area at 3dpf (**I**) and 5dpf (**J**) compared to controls.

The transgenic line *Tg(−4.9sox10:EGFP)* (Carney et al., 2006) that labels pre-migratory and migrating CNCCs was used to examine these cells because of their large contributions to craniofacial bone and cartilage. Early in development (24hpf, **Fig. 6A**) CNCC numbers and migration were unaffected in the crispants and morphants when compared to controls (p = 0.27). However, by 48hpf, significant differences were observed in the number and migration pattern of CNCCs in *fos* crispants and morphants (p < 0.0001) (**Fig. 6A, B**). At 3dpf, crispants and morphants continued to show reduced CNCC derivatives compared to both UICs and MMs and these differences were more pronounced at 5dpf, with significant decrease of GFP fluorescence in crispants/morphants compared to UIC controls (p < 0.0001 for both, **Fig. 6A, B**). Activated caspase-3 showed a dramatic increase in expression around the eyes and in the midline in both crispants and morphants suggesting high, localized cell death between 24hpf and 2dpf (**Fig. 6C**). While MOs are known to cause an increase in cell death (Boer, Jette, & Stewart, 2016), the CNCC apoptosis was also observed in the F0 crispant larvae, suggesting this precocious death is promoted by a lack of *fos*. Together, these results demonstrated that absence/knockdown of *fos* compromised the survival of critical subpopulations of CNCCs and contributed to the phenotypic anomalies observed later in development.

**Figure 6.**
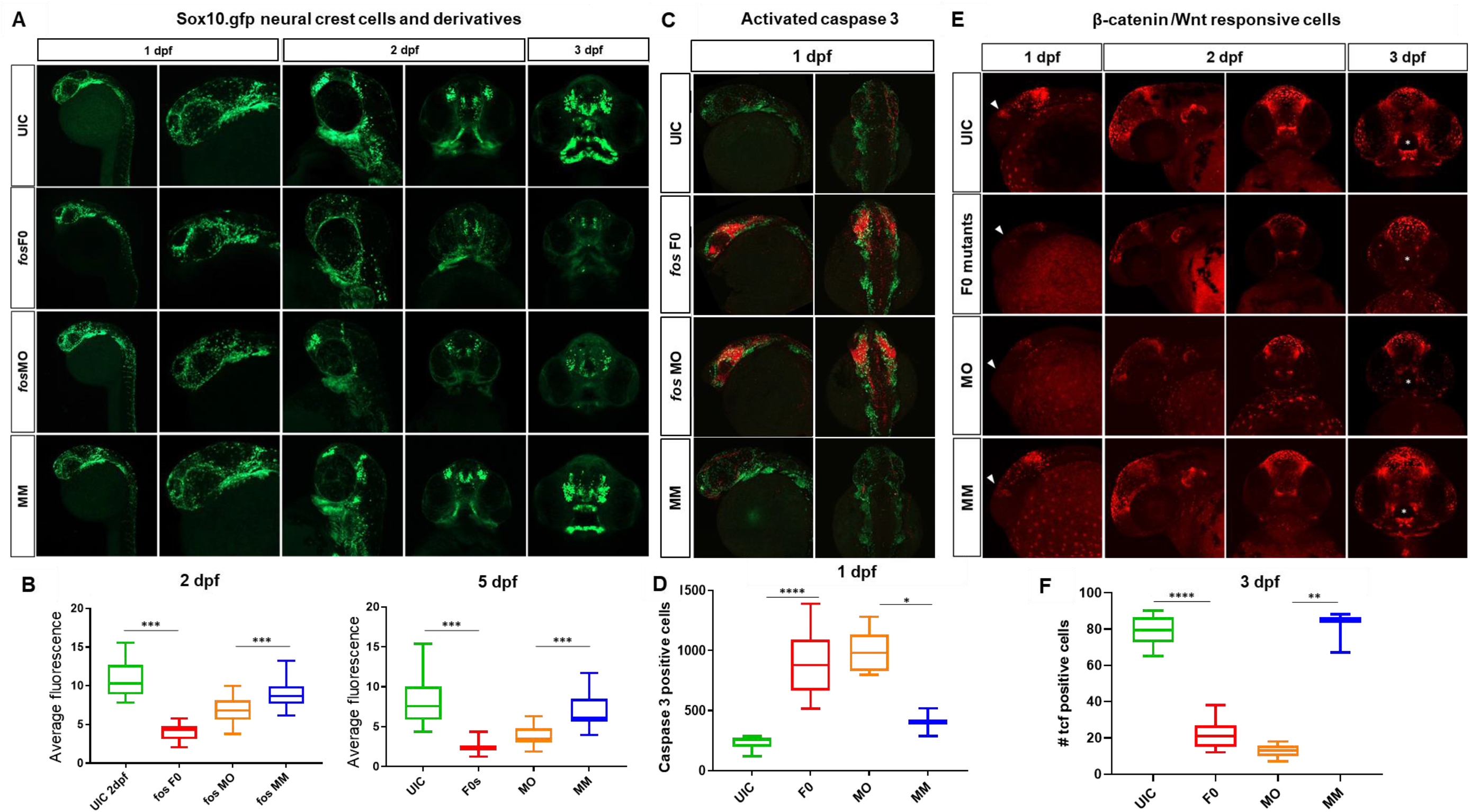
Craniofacial cell populations are significantly affected in *fos* crispant and morphant zebrafish embryos. **A**) Time course evaluation of CNCCs shown at 24hpf, where sox10 fluorescence was similar in all 4 groups, but at 48 hpf significant differences in NCC populations and migration were observed in the *fos* crispants and morphants and these differences persisted to 5 dpf in NCC derived mesenchyme as observed in rostral views of embryos. **B**) Quantification of average fluorescence showed significant reductions at both 48hpf and 5dpf for crispants and morphants. **C**) *Fos* crispants/morphants showed increased apoptosis in the head region at 24 hpf, shown by activated caspase 3 immunolabeling, where affected CNCC populations around the eye and optic stalk will give rise to the ethmoid plate and oral ectoderm. **D**) The increase in positively stained cells was significantly increased in crispants and morphants compared to controls. **E)** Mutation and knockdown of *fos* in a Wnt reporter line showed a reduction in β-catenin signaling starting at 1 dpf (triangles), when a group of cells observed in controls was largely missing in the mutants/morphants and persisting to 3 dpf, when reporter expression in cells of the upper and lower lip was severely reduced in fos mutants and morphants (white asterisks) **F**) 3D quantification of the oral cavity region supported the significant reduction in Wnt-responsive cell number in these structures. UIC = uninjected control, MO = *fos* morpholino, MM = mismatch control morpholino. * p < 0.05, ** p < 0.01, *** p < 0.001, **** p < 0.0001.

Interactions between the mesenchyme and epithelial cells are known to induce Wnt signaling centers that control cell fates and behaviors in the surrounding tissues (Jussila & Thesleff, 2012). Based on the changes observed in CNCC and epithelial cells, a reporter zebrafish line that serves as a biosensor for Wnt signaling (*Tg(7xTCF-Xla.Siam:GFP)*) was evaluated for changes in β-catenin mediated Wnt activity (Moro et al., 2012). A general reduction in WNT-responsive cells, in both *fos* crispants and morphants, was identified in the head starting at 1 dpf and 2 dpf (p < 0.01, **Fig. 6, Supplemental Fig. 13**). At 3 dpf, we observed a group of WNT responsive cells that were restricted to the upper and lower lip. Intriguingly, a dramatic reduction of these Wnt responsive cells was observed in both *fos* crispants and morphants (**Fig 6**, asterisks). Quantification of these cells in 3D analysis using Imaris supported the significantly reduced number of reporter positive cells in the oral cavity region (p = 4.08E-12 and p = 0.007, crispants compared to UIC and morphants compared to MM controls, respectively). By 5dpf, β-catenin active cells had recovered around the oral cavity in crispants/morphants, however the organization of the cells followed the abnormal mouth shape. Together, these results suggested that Wnt responsive cells were reduced at early developmental time points and severely disrupted around the mouth at 3 dpf, implicating that absence or reduction of *fos* impacted the ability of these cells to properly pattern the oral cavity.

## DISCUSSION

*Fos* plays a role in embryonic and craniofacial development (Alfaqeeh et al., 2015; Durchdewald et al., 2009; Velazquez et al., 2015; Wagner, 2002) and was previously identified in a network of differentially expressed genes in a zebrafish model of orofacial clefts and confirmed in a human family-based association study (Chiquet et al., 2018). In this study, we sought to characterize the role of Fos in orofacial morphogenesis and oral cavity formation. Using both *fos* crispant and morphant embryos, we showed that that absence or reduction of *fos* in developing zebrafish led to an abnormal craniofacial morphology with a distinctive abnormal ovoid oral cavity shape, suggesting a key role in orofacial morphogenesis. Deep phenotyping using geometric morphometric analyses showed that midface dimensions and oral cavity shape were the facial features most affected by *fos* perturbation. Further, perioral epithelial cells had altered morphologies and cranial neural crest cells (CNCCs) were reduced in number. These changes corresponded with a reduction in Wnt-responsive cells around the oral cavity, suggesting that *fos* plays an important regulatory role in guiding the interactions between the epithelial and mesenchymal cell populations that induce WNT signaling during oral cavity development. Together, our findings provide a better understanding of the contributions of this gene in vertebrate craniofacial development and suggest that variation in *FOS* expression can alter facial dimensions during development and potentially play an etiologic role in orofacial clefting.

Cell migration and movements during craniofacial development facilitate interactions between different cell and tissue types, and their environment (Murillo-Rincon & Kaucka, 2020). These interactions are critical to create spatially defined sources of morphogenetic signals (e.g., WNTs, FGFs, BMP, Hedgehog), commonly referred to as signaling centers or organizers (Martinez Arias & Steventon, 2018; Murillo-Rincon & Kaucka, 2020; Perrimon, Pitsouli, & Shilo, 2012). Both the initiation of the signaling centers in the developing head and the expression levels of inductive signals can be modified by several mechanisms, such as 1) changes in gene expression that perturb survival or migration of specific cell types and 2) the timing of the cell/tissue interactions (Perrimon et al., 2012).

Although CNCC death can be associated with morpholino oligonucleotide use (Boer et al., 2016), the marked upregulation of apoptosis of CNCC in the head and around the eyes was also identified in the crispants, but not in mismatch morpholino injected embryos. These results suggest that the observed apoptosis is not solely due to morpholino-related toxicity. The CNCC precursors in these regions have been shown to migrate rostrally and caudally to reach the oral ectoderm and ethmoid plate (Eberhart et al., 2008) and the observed aberrant apoptosis likely led to the reduction of CNCCs, resulting in abnormal arches and anomalous jaw skeletal elements. Indeed, *fos* knockdown caused increased apoptosis in osteosarcoma cells (Wang et al., 2017), supporting the relationship between reduced levels of fos protein and increased apoptosis observed in our crispant and morphant zebrafish models. In *C. elegans, fos* is also a cell autonomous regulator of cell invasion, a process critical for many cell types and also NCCs, as they need to invade through extracellular matrix, mesoderm and migrating endothelial cells to reach their destination making their motility and invasive ability crucial (McLennan et al., 2020; Sherwood, Butler, Kramer, & Sternberg, 2005). Overall, perturbation of zebrafish *fos* expression affected the survival and migration of a specific subset of NCCs that populate the oral ectoderm and ethmoid plate, suggesting that one of the mechanisms responsible for the abnormal orofacial phenotype is the early disruption of NCCs.

Epithelial-mesenchymal interactions play a crucial role during the development of several organs and tissues and are mediated by cell-cell contact, or the diffusion of soluble factors such as Wnts (Ribatti & Santoiemma, 2014). During tooth development, regional signals from the ectoderm induce molecular changes in the cranial neural crest-derived ectomesenchyme (Jussila & Thesleff, 2012). The activation of β-catenin signaling is required in the oral epithelium for the induction of placodes and formation of enamel knots; this signaling is also necessary in the odontogenic mesenchyme beyond the bud stage (Chen, Lan, Baek, Gao, & Jiang, 2009; Jussila & Thesleff, 2012). Both *fos* crispants and morphants showed a dramatic reduction of Wnt-responsive cells around the oral cavity at 3dpf, whereas the brain was less affected, indicating that Wnt signaling was perturbed in this region. Consistent with disrupted inductive signaling, *fos* crispants/morphants showed abnormal tooth development, with marked delays in tooth formation and anomalies in morphology and number of teeth. Even though the process of tooth eruption in mice is different from pharyngeal tooth attachment to CB5 in zebrafish, these results are consistent with observations of smaller teeth in *Fos* null mice that fail to erupt and form roots (Alfaqeeh et al., 2015; Van der Heyden & Huysseune, 2000).

The periderm serves important functions in craniofacial structures that fuse during development, such as the oral cavity and the palate (Hammond, Dixon, & Dixon, 2019). In mammalian models, abnormal periderm development leads to epithelial adhesions and orofacial clefts, while failure of periderm removal from fusing palatal shelves can contribute to cleft palate (Hammond et al., 2019; Richardson et al., 2014). Periderm anomalies were observed starting at 3dpf in the perioral region, suggesting that loss of *fos* could also be contributing to peridermal abnormalities linked to abnormal orofacial development. Future experiments will identify how *fos* regulates the molecular mechanisms in periderm cells. Interestingly, our results showed that at 3 dpf, CNCC derivatives were greatly reduced in the upper and lower jaws, at the same time that oral periderm cells showed altered morphologies and arrangement, indicating that perturbation of *fos* interrupted both epithelial and mesenchymal cell populations around the developing oral cavity. This specific effect on cells around the oral cavity corresponds with the mouth and midface shape anomalies detected at 5dpf, suggesting a disruption to critical inductive signals that regulate orofacial morphology. This could result from either missing β-catenin active cells or inability of these cells to activate β-catenin signaling and warrants further studies. These findings suggest that *fos* plays an important inductive role in regulating interactions between epithelial and mesenchymal cell populations and its absence affects both cells around the oral cavity and midface and mineralized tissue like bones and teeth.

Additionally, Wnt/β-catenin signaling delineates areas of rapid growth in the facial prominences in murine models, and both genetic and pharmacologic perturbation of this signaling pathway results in altered craniofacial dimensions during embryonic development (Brugmann et al., 2007; He & Chen, 2012). The use of geometric morphometrics allowed identification of regional alterations to dimensions such as mouth height, olfactory to mouth angles and olfactory distance, and identified significant shape changes to the mouth and midface, strongly supporting the role of *fos* in morphogenesis of these structures (Maili et al., 2022). Together with the reduced number of Wnt responsive cells in the perioral region, these findings suggest insufficient growth in these tissues when *fos* is absent or reduced. Interestingly, a correlation between the dimensions of the maxillae and nasal pits and susceptibility for developing cleft lip and palate has been reported in mice (Green et al., 2015; Parsons et al., 2008; Young et al., 2007). Additionally, human morphometric studies have shown that there are facial differences in the upper lip, philtrum and nasolabial angles in individuals predisposed to orofacial clefts as well as their unaffected relatives (El Sergani et al., 2020; Fraser & Pashayan, 1970; Weinberg, Maher, & Marazita, 2006; Weinberg et al., 2009; Weinberg, Neiswanger, et al., 2006). Together with the findings that variation in *FOS* is associated with NSCLP, these results suggest that altered *fos* expression affects the growth and shape of the oral cavity and midface regions and potentially plays an etiologic role in the development of an orofacial cleft.

Crispants, morphants and pharmacologic inhibition of the transcription factor complex AP-1 produced similar orofacial anomalies, supporting our findings. Stable *fos* mutants (F1, F2, F3 generations) had no phenotype even though they had 1.5 kb deletions and absent *fos* mRNA expression and thus were not useful for these studies. Similar results from multiple studies suggest that this might be due to genetic compensation unique to Crispr/Cas9 mutagenesis methods (Buglo et al., 2020; Rossi et al., 2015). Increased expression of *fosl1a, fosb* and *fosl2* was observed in the *fosab* mutants (Sup. Fig 3). Additionally, injection of the guide RNAs in *fosab* mutant embryos did not produce a phenotype (Sup. Fig 4). These data support the idea that compensatory mechanisms by similar gene family members may be responsible for buffering the phenotype.

In conclusion, this study demonstrates that *fos* is critical for zebrafish craniofacial development and *fos* perturbation affects multiple craniofacial tissues resulting in abnormal orofacial structures. Loss of *fos* has a profound effect on facial progenitor cells, suggesting that it is important for epithelial and mesenchymal cell populations and regulates interactions between them that are crucial for proper oral cavity morphogenesis. Although cleft lip was not seen in our zebrafish embryos, severe orofacial anomalies were observed. This result may be related to species-specific differences in mouth development (Soukup, Horacek, & Cerny, 2013). Based on the current work as well as results suggesting a role for *FOS* in NSCLP (Chiquet et al., 2018), future investigations in zebrafish and humans should target identification of other genes and pathways by which *fos* regulates craniofacial development.

## MATERIALS AND METHODS

### Zebrafish care and husbandry

Zebrafish (*D. rerio*) were housed and maintained at 28^0^C as described previously (Westerfield, 1993). All work involving the use of animals was performed with approval of the UTHealth Animal Welfare Committee (AWC-20-0052).

### Morpholino, mRNA and CRISPR/Cas9 injections

Zebrafish *fos* antisense morpholinos (Fos MO: GCGTTAAGGCTGGTAAACATCATCC) targeting the ATG start site in exon 1 and a mismatched control (Fos MM: GaGTTAAcGCTcGTAAAaATCATaC, mismatch in small caps) were designed by GeneTools (Philomath, OR). Morpholinos were suspended in MilliQ water to a stock concentration of 16.67mg/mL or 2mM. Injections of morpholinos were diluted to 0.5 ng/uL to 3ng/nL in Danieu buffer. A plasmid containing the full-length human *FOS* cDNA (NM_005252.4) was purchased from Addgene (Plasmid #59140). The full-length *FOS* cDNA was cloned into the pCS2 vector and *FOS* mRNA was generated using the mMessage mMachine Sp6 kit (Ambion, Austin, TX). mRNA was resuspended in nuclease-free water to a stock concentration of 2ng/nL and diluted to 0.5ng/nL in 0.1M KCl for injections. Fos F0 crispants were created using IDT Alt-R™ CRISPR-Cas9 System (Coralville, IA). Two Crispr RNAs (crRNAs) specific to Fos gene (*fos* crRNA1: CGAGCAAGGAAATACAAGAC and *fos* crRNA2: GGTTGGGGAATTCAAGGAGT) were hybridized separately with trans activating crispr RNA (tracrRNA) to form a functional gRNA complex. 60uM of each gRNA was incubated with 5ug/ul of Cas9 protein (Alt-R^®^ S.p. Cas9 Nuclease, IDT) for 10 minutes at 37^0^C to generate the ribonucleoprotein (RNP) complex. Equimolar amounts of the two RNPs were mixed and injected into the zebrafish embryos. For all zebrafish injections, one-cell embryos were injected with 1nL of MO, mRNA or RNPs.

### Zebrafish Facial Analytics based on Coordinate Extrapolate system (zFACE)

Embryos were fixed in 4% paraformaldehyde (PFA) (Sigma) in 1X phosphate buffered saline with Triton X-100 (PBST) at room temperature for 4 hours and stained with 0.2mg/L DAPI (Life Technologies) for 30 minutes at room temperature. Embryos were mounted rostrally in 1% low-melt agarose (Research Products International, Mount Prospect, IL) and imaged with Zeiss LSM 800 Confocal Microscope (Thornwood, NJ). Twenty-six anatomic landmarks including eyes, olfactory pits, neuromasts and mouth were identified from the confocal images and measurements between these landmarks were calculated to extract phenotypic features and understand which anatomical structures were altered as a result of *fos* knockdown. ANOVA and Tukey’s test for multiple comparisons was applied or each measurement and Bonferroni correction for 39 measurements was applied to determine statistical significance. GraphPad Prism 9.0.0 was used to plot and visualize the data.

### Multivariate analysis of zFACE measurements and landmarks

Dimensionality reduction was performed using Principal Component Analysis (PCA) in StataIC 14 (StataCorp. 2015). Components with an eigenvalue of greater than or equal to 1 (following the Kaiser-Guttman method) were retained for analysis. Promax rotation, which accounts for correlations between the different factors (zFACE measurements) was used because a high correlation was observed/ expected between features calculated using shared landmarks. Principal component (PC) scores were predicted and logistic regression models were utilized to regress morphant/crispant status by PC scores.

Additionally, to focus on facial shape and remove variation due to size, position, or rotation, the 2D landmark data points from the 26 zFACE coordinates were uploaded into MorphoJ version 1.07A and principal axes Procrustes superimposition was performed (Klingenberg, 2011). After Procrustes transformation, PCA was used to examine general shape variation in the combined groups. Additionally, discriminant function analysis (DFA), which maximizes the between-group variance relative to within-group variance, was used to pinpoint shape in *fos* morphants/crispants compared to controls. Mean Procrustes distance with 10,000 permutations was used to statistically test shape differences between control and experimental groups.

### Skeletal staining

Alcian blue (Anatech LTD) and alizarin red (Sigma-Aldrich) staining was performed using standard techniques (Kimmel and Trammell 1981) to visualize the bone and cartilage structures. Briefly, 5-8dpf embryos were collected and fixed in 2% PFA/1X PBXT for 1 h at room temperature and stored in methanol over night at −20°C. After removing methanol, embryos were incubated in 0.04% alcian blue solution (100 mmol/L Tris, pH 7.5, 10 mmol/L MgCl2, 64% ethanol) overnight at room temperature. They were destained in 3% H2O2/0.5% KOH for 10 min at room temperature and then stained in 0.02% alizarin red solution (100 mmol/L Tris, pH 7.5, 25% glycerol) for 30 min at room temperature. Embryos were then destained in 50% glycerol/0.1% KOH for 30 min and stored in 50% glycerol. Imaging was performed using the LAS Montage Module (Leica). For visualizing teeth, a 0.5% Alizarin Red solution was used to stain fixed sox10.gfp reporter embryos for 1 hour.

### Fluorescent signal and cell measurements

Periderm cells were measured in Zen software (Zeiss, Thornwood, NJ) using the contour graphics tool. Midface and perioral cells were analyzed separately due to already existing differences in their size. In each individual embryo, 10 cells were measured from each region and at least 10 individuals were analyzed from each group.

Average fluorescence signal was measured in ImageJ (Schneider, Rasband, & Eliceiri, 2012). Wnt-responsive cells were counted using the Spots detection tool and automatic detection settings in Imaris software (Bitplane, Oxford Instruments) utilizing default settings with at least 3 animals analyzed for each condition.

### Caspase assay

Embryos were fixed at 1dpf in 2% PFA/1XPBS overnight at 4°C. After washing, they were incubated with block solution (1% DMSO, 2mg/ml BSA, 0.5% Triton X-100, 10% inactivated goat serum in PBS) for 2 hours at room temperature, washed and then incubated with 1:700 dilution of Rabbit Activated Caspase 3 (BD Biosciences, Franklin Lakes, NJ) in block solution overnight at 4°C. 1:150 dilution of Donkey Anti-Rabbit 647 (Molecular Probes, Eugene, OR) was used for secondary antibody staining. Washed embryos were mounted in 1% low-melt agarose for confocal imaging.

### HCR assay

HCR assay was performed using Molecular Instruments Multiplexed v3.0 protocol and reagents (Molecular Instruments, Los Angeles, CA). Briefly, embryos were fixed at different stages in 2% PFA/1XPBS over night at 4°C. They were pre-hybridized with probe hybridization buffer for 30 minutes at 37°C, then incubated with 4nM of the probe (*fosab* HCR probe set B2 Alexa Fluor 546) in probe hybridization buffer overnight at 37°C. Embryos were washed with 5X SSCT and incubated with amplification buffer for 30 minutes at room temperature. 30 pmol of hairpins h1 and h2 corresponding to the probe were snap cooled (heated at 95C for 90 seconds and cooled to room temperature in a dark drawer), diluted in 500uL of amplification buffer and added to the embryos for overnight incubation in the dark at room temperature. Washed embryos were mounted in 1% low-melt agarose for confocal imaging.

### Quantitative PCR

Total RNA was extracted from a group of 5 embryos using Trizol (Invitrogen) as described previously (Chiquet et al., 2018). Further purification was done using RNeasy Mini Kit (Qiagen). The purified RNA was reverse transcribed to cDNA using Quantitect™ Reverse Transcription Kit (Qiagen). Gene expression of different *fos* family genes in zebrafish was analyzed using Quantitect™ Primer Assays (Qiagen) for each individual gene – *fosaa, fosab, fosb, fosl1a, fosl1b* and *fosl2* – using beta-actin (*actb1*) as an endogenous control. All samples were run in triplicate and compared using Student’s t-tests.

### Pharmacological treatments

AP-1 inhibitor SR 11302 (Tocris Bio-Techne, Minneapolis, MN) was dissolved in DMSO to prepare a stock solution. Varying concentrations of drug (1uM, 5uM and 10uM) were added to embryo dishes starting at either 1 or 2 dpf with daily media changes until collection at 5 dpf. The controls included treatment with the same volume of DMSO and untreated dishes.

## Acknowledgements

We thank Christian Urbina for morphometric data analysis assistance.

## Data Availability

The data that support the findings of this study are available from the corresponding author upon reasonable request.

## Author contributions statement

L.M., B.T., A.L., G.T.E. and J.T.H conceptualized the project and methodology. L.M., B.T., Q.Y., and S.M. performed all the experiments and subsequent analyses. S.S.H. provided assistance with statistical analysis. L.M and B.T. wrote the manuscript and A.L., G.T.E. and J.T.H. edited the manuscript.

